# Social context and dopamine signaling converge in the mushroom body to drive impulsivity

**DOI:** 10.1101/2025.02.21.639508

**Authors:** Paul Rafael Sabandal, Young-Cho Kim, John Martin Sabandal, Kyung-An Han

## Abstract

Organisms adapt their behaviors flexibly in response to various internal and environmental factors. However, how and where these factors converge in the brain to alter behavior is not well understood. In this study, we examine how social context interacts with dopamine activity to influence inhibitory control in *Drosophila*. We found that, regardless of social context—whether isolated or in groups—wild-type flies consistently showed strong movement suppression in a go/no-go task that measures action restraint. In contrast, flies with enhanced dopamine activity suppressed their movements when tested alone or with potential mates but exhibited impulsive behaviors when exposed to same-sex peers. This social-context-dependent impulsivity was shown to rely on dopamine-D1 receptor-cAMP signaling in mushroom body (MB) neurons. Remarkably, activating the MB was sufficient to induce impulsivity, even without dopamine input or a social context. Our findings highlight MB as a critical hub where social context and dopamine signaling converge to regulate impulsive behavior in *Drosophila*.

**Signficance statement:** This study demonstrates that impulsivity results from the interplay between elevated dopamine levels and social context, rather than dopamine alone, with the mushroom body (MB) serving as a key neural hub for integrating these signals in *Drosophila*. Social stimuli, such as the presence of same-sex peers, disrupt inhibitory control in a context-dependent manner, highlighting the importance of multimodal sensory inputs and MB activity. These findings challenge the isolation-focused approach in traditional impulsivity research and underscore the need to account for social influences when investigating cognitive processes and disorders like ADHD, autism, and substance use, where social settings often amplify symptoms.

**Classification:** Genetics / Neuroscience

## Introduction

Inhibitory control enables individuals to stop pre-planned or ongoing actions or thoughts that are inappropriate at a given context. Effective inhibitory control allows us to halt crossing a street when an unexpected drunk driver speeds by or to suppress the impulse to accelerate when a traffic light turns yellow. Similarly, sugar gliders refrain from foraging during storms, and toadfish stop calling when they detect predator signals ^1,2^. This cognitive process is essential for performing daily activities ^3,4^. Dysfunctional inhibitory control leads to impulsivity—a tendency to act without forethought ^3–5^, compromising an individual’s fitness and survival. Impulsivity is a core feature of several neurological and psychiatric disorders, including attention deficit hyperactivity disorder (ADHD), autism spectrum disorder, obsessive-compulsive disorder, bipolar disorder, post-traumatic stress disorder, sleep disorders, and substance use disorders ^3–8^.

The neural structures and neurotransmitters involved in inhibitory control and impulsivity have been extensively studied in both humans and animal models ^3–5^. For instance, individuals with focal lesions in the frontal lobe poorly perform the go/no-go task, a test that assesses action restraint and motor impulsivity ^3,9,10^. Additionally, altered dopamine signaling has been shown to impact cognitive and motor impulsivity ^5,9^. Positron emission tomography (PET) imaging studies have demonstrated that dopamine receptor availability in the cortex changes during a stop-signal task (SST), a test measuring action cancellation. These changes are positively correlated with dopamine release in brain regions associated with inhibitory control ^11^. Furthermore, polymorphisms in genes encoding dopamine transporter (DAT) and D1, D2, and D4 receptors are associated with altered inhibitory control and frontal lobe activity ^12–14^. These studies establish a strong link between dopamine neurotransmission and impulsivity. However, the underlying mechanisms of this relationship remain poorly understood.

Impulsivity is multifaceted, influenced by both genetic factors (e.g., dopamine-related genes) and non-genetic factors, such as drug use ^15^, aging ^16–18^, and early life experiences ^19,20^. Notably, there is limited understanding of how genetic and non-genetic risk factors interact to influence impulsivity. Additionally, most studies on inhibitory control and impulsivity in humans and animal models are conducted in socially isolated settings, despite the fact that most organisms live in dynamic social environments. The impact of social context on inhibitory control and impulsivity remains largely unexplored.

In this report, we address these gaps using the *Drosophila* model. We demonstrate that dopamine interacts with social context to modulate behavioral responses in a fly-version of the go/no-go task, which measures action restraint. Flies with elevated dopamine levels show robust movement suppression when tested in isolation but fail to exhibit the same control in the presence of same-sex peers. Moreover, we identify the dopamine-D1 receptor-cAMP signaling pathway in the mushroom body (MB) γ neurons as a key mediator of impulsive behavior in group settings.

## Methods

### Fly stocks and culture

The wild-type strain used in the study is *Canton-S* (*CS*). We previously described *fumin* (*fmn*) ^21,22^, *dumb*^2^ (*dumb*) ^23^, *der* ^24^, *damb* ^25^, *dd2r* ^26^, all of which are in the *CS* background, as well as *MB247-GAL4* ^23,27^ from Dr. Waddell (University of Oxford, Oxford, UK); *NP1131-GAL4* ^28^ from Dr. Dubnau (Stony Book University School of Medicine, Stony Brook, NY); *TH-GAL4* ^29^, *UAS-dDA1* ^23^, and *c547-GAL4* ^30^. *DDC-GAL4* (stock no. 7009) ^31^*, TRH-GAL4* (38388), *TDC2-GAL4* (9313), *c739-GAL4* (7362) ^32^, *c305a-GAL* (30829) ^32^*, elav-GAL4* (8765) ^23^*, Orco-GAL4* (26818) ^33^, *Gr32a-GAL4* (57622) ^34,35^, *Gr63a-GAL4* (9942) ^36^, *OK107-GAL4* (854) ^27^, *30Y-GAL4* (30818) ^32^, *NPF-GAL4* (25682) ^37^, *c232-GAL4* (30828) ^32^, *c205-GAL4* (30826) ^32^, *MB058B-GAL4* (68278), *MB304B-GAL4* (68367), *MB308B-GAL4* (68312), *MB630B-GAL4* (68334), *MB320C-GAL4* (68253), *MB438B-GAL4* (68326), *MB504B-GAL4* (68329), *MB060B-GAL4* (68279), *MB065B-GAL4* (68281), *MB099C-GAL4* (68290), *MB296B-GAL4* (68308), *MB043C-GAL4* (68363), *MB043B-GAL4* (68304), *MB299B-GAL4* (68310), *MB194B-GAL4* (68269), *MB063B-GAL4* (68248), *MB213B-GAL4* (68273), *MB025B-GAL4* (68299), *MB301B-GAL4* (68311), *MB056B-GAL4* (68276), *MB032B-GAL4* (68302), *MB195B-GAL4* (68270), *MB441B-GAL4* (68251), *MB188B-GAL4* (68268), *MB042B-GAL4* (68303), *MB196B-GAL4* (68271), *MB047B-GAL4* (68364), *MB312B-GAL4* (68314), *MB312C-GAL4* (68252), *MB316B-GAL4* (68317), *MB109B-GAL4* (68261), *MB313C-GAL4* (68315), *MB315C-GAL4* (68316), *MB298B-GAL4* (68309), *MB210B-GAL4* (68272), *MB434B-GAL4* (68325), *MB433B-GAL4* (68324), *MB399B-GAL4* (68329), *MB002B-GAL4* (68305), *MB074C-GAL4* (68282), *MB310C-GAL4* (68313), *MB112C-GAL4* (68263), *MB083C-GAL4* (68287), *MB110C-GAL4* (68262), *MB057B-GAL4* (68277), *MB051B-GAL4* (68275), *MB051C-GAL4* (68249*), MB077B-GAL4* (68283), *MB077C-GAL4* (68284), *MB050B-GAL4* (68365), *MB549C-GAL4* (68373), *MB080C-GAL4* (68285), *MB062B-GAL4* (68280), *MB093C-GAL4* (68289), *MB082C-GAL4* (68286), *MB543B-GAL4* (68335), *MB018B-GAL4* (68296), *MB542B-GAL4* (68372), *MB242A-GAL4* (68307) ^38^, *UAS-TrpA1* (26264) ^39^, *UAS-shi^ts^* (44222) ^40^, and *UAS-Epac1-camps* (25407, 25408)^41^ were obtained from the Bloomington Drosophila Stock Center (Bloomington, IN); *TH-C’* and *TH-D1* ^42^ from Dr. Wu (Johns Hopkins University, Baltimore, MD); *pBDP-GAL4* ^43^ from Dr. Anderson (California Institute of Technology, Pasadena, CA); *R58E02-GAL4* ^44^ from Dr. Tomchik (Scripps Research Institute, Jupiter, FL); the *rutabaga*^2080^ allele ^45^ and the *dunce*^1^ allele^46^ from Dr. Davis (Scripps Research Institute, Jupiter, FL).

*TH,DDC-GAL4* was made by recombining *TH-GAL4* and *DDC-GAL4* on the third chromosome and used in the experiments manipulating all dopamine neurons since it labels more PAM neurons than *TH-GAL4*. The open reading frame of *Drosophila* DAT was cloned under UAS in the gateway vector pTW ^47^ and transgenic *UAS-DAT* lines were generated by germ line transformation in *w*^1118^ embryos. Germ-line transformed lines were outcrossed with Cantonized *w*^1118^ for six generations and the transgenes in the third chromosome were placed in the *fmn* genetic background.

All fly stocks were reared on a standard cornmeal/agar medium at 25° C with 50% relative humidity under the 12 h light/12 h dark cycle. Flies were collected under CO_2_ within two days after eclosion and used when they were 4 – 5 days old. To prepare dead flies, *CS* flies were frozen and brought back to room temperature before use. Decapitated flies were used within 15 min after their heads were cut off.

### Go/no-go test

The go/no-go test was done as previously described ^18^. Briefly, flies were placed in a rectangular plexiglass chamber (60 mm L X 60 mm W X 15mm H) connected to filtered air and acclimated to the chamber for 10 min. The 10 L/min airflow or recorded wasp sound was delivered to the chamber for 10 min. All experiments unless otherwise stated were performed with the airflow. The chamber was video recorded to monitor fly movements before and after airflow. Videos were analyzed using the Viewer3 tracking software (BiObserve Technologies, Bonn, Germany), which allows tracking and measuring the average speed in mm/sec per fly. Raw data were transported to Excel (Microsoft, Redmond, WA) and the number of the movements exceeding 60 mm/sec that we defined as loss of inhibition events (LIE) was scored per fly per min. The highest LIE values were used for comparison. Genotypes and flies with drug treatments were blinded to experimenters.

For activation or inhibition of neuronal activity, the flies carrying selected *GAL4* and *UAS-TrpA1* or *UAS-shi^ts^*, respectively, were placed in a heat box at either 25° C (control) or 30° C (activation or inhibition) and subjected to the go/no-go test. A heat box (internal dimensions: 360 mm L x 170 mm W x 170 mm H) was made from a hybridization oven (Stovall, Greensboro, NC) by removing rotating racks. The webcam LifeCam Studio (Microsoft, Redmond, WA) installed in the heat box was used for video recording.

### Pharmacological treatment

The 3 days-old flies were housed in the food vial containing different concentrations (0, 1, 5, 10 and 20 mg/ml) of either D1 receptor antagonist SCH-23390 hydrochloride (D054, Sigma-Aldrich, Saint Louis, MO) or D2 receptor antagonist Eticlopride hydrochloride (E101, Sigma-Aldrich) for 24 h. Both D1 and D2 antagonists have been shown to be effective on the fly receptors ^48^. Ascorbic acid (0.25 mg/ml, A92902, Sigma-Aldrich) and green food color (McCormick & Company, Inc., Sparks, MD) were added to minimize oxidation of the drugs and to visualize fly abdomens to monitor intake, respectively.

For ablation of MB neurons, first instar larvae were collected within 6 h after hatching and fed with yeast paste containing 20 mg/ml hydroxyurea (H8627, Sigma-Aldrich) for 4 h ^49^. The larvae were rinsed in distilled water and then transferred to fresh food. The 4-to 5-day old flies were collected and used for go/no-go tests where untreated flies served as a control.

### cAMP imaging

The *fmn* or control flies expressing the Epac1-camps ^41^ in the MB (through *OK107-GAL4*) were used for imaging. Fly brains were dissected in ice-cold hemolymph-like saline (HL3; 70 mM NaCl, 5 mM KCl, CaCl2 1.5 mM, 20 mM MgCl2, 10 mM NaHCO3, 5 mM Trehalose, 115 mM Sucrose and 5 mM Hepes, pH 7.1) within a couple of minutes, placed on a microscope glass containing either HL3 (control) or 100 µM dopamine in HL3 (treated) and then covered with a cover slip. Two layers of the scotch tape (3M Company, St. Paul, MN) were used to create space between a microscope glass and a cover slip. Imaging was done using the Zeiss LSM700 confocal microscope (Zeiss, Thornwood, NY) within 2 min after dissection. Excitation of Epac1 was done at 440 nm and emission at 480 nm for CFP and 540 nm for YFP. Optical sections were made every 2 μm with the 512 x 512 pixel resolution. Peak cAMP response in the medial lobes per hemisphere was measured by the inverse FRET ratio, which is |ΔR/R_0_|, where R = CFP/YFP and ΔR = R_treated_ – R_non-treated_ ^41,50^.

### Statistical analysis

Statistical analyses were performed using Minitab 20 (Minitab, State College, PA). All data are reported as mean + or ± standard error of mean (SEM). Normality was determined by the Anderson Darling goodness-of-fit test. The normally distributed data of two groups were analyzed by a two-tailed Student’s *t*-test. The data with three or more groups were analyzed by General Linear Model (GLM) with either *post hoc* Dunnett’s or Tukey tests. Non-normally distributed data sets were analyzed by Mann-Whitney and Kruskal-Wallis tests.

## Results

### Hyper-dopamine activity in a group, but not a solitary, setting leads to impulsivity

We previously reported that aging is a key risk factor for impulsivity with cholinergic signaling playing a major role in this process ^18^. Building on this, we investigated whether social context influences inhibitory control. To test this, we exposed single flies and groups of varying sizes (2, 4, 8 or 13) to a GNG assay. In the absence of strong airflow, wild-type *Canton-S* (*CS*) flies moved freely (Go phase in Figures 1A and 1B). When a strong airflow (10 L/min) was introduced (No-Go phase), the flies stopped moving and remained stationary for the duration of the No-Go phase with occasional, sporadic flights (Figures 1A and 1B). The frequency of these sporadic flights did not increase with the number of flies in the chamber (Figure 1C), suggesting that social context does not affect inhibitory control of wild-type flies.

**Figure 1.**
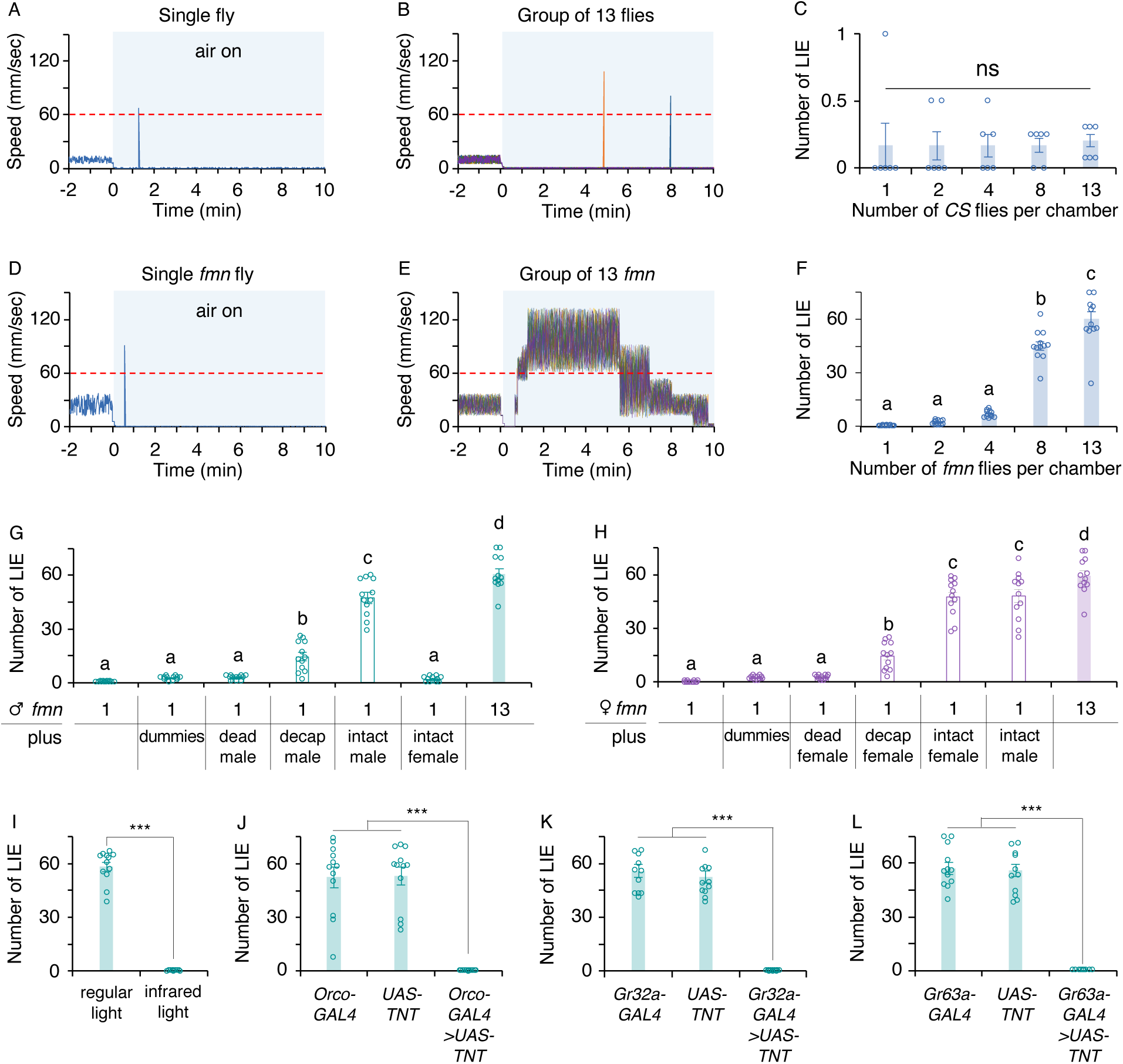
The mere presence of others along with hyper-dopamine triggers impulsivity, and this process involves multiple sensory cues. The mere presence of others does not affect impulsivity. **A-B.** Representative movement speed traces of a chamber containing a single wild-type *Canton-S* (*CS*; **A**) and of a chamber containing 13 *CS* flies (**B**) under the Go/No-Go test (GNG). The white area indicates Go phase where the airflow is off, the shaded blue area denotes No-Go phase where the 10 L/min airflow is on, and the traces exceeding the dotted red line on the 60 mm/sec mark represent a loss of inhibition event (LIE). The *CS* exhibited exploratory behavior in the Go phase but stopped moving upon exposure to airflow and continued to show movement suppression in the No-Go phase with sporadic LIEs (**A-B**). **C.** Manual counting of LIEs per fly per min revealed that the number of *CS* flies in chamber does not impact impulsivity. (1, 2, 4, 8, versus 13; Kruskal-Wallis test, *p* = 0.481, *n* = 6). The dopamine transporter mutant *fumin* (*fmn)* exhibit heightened impulsive flying under the GNG when tested in a group. **D-E.** Representative movement speed traces of a chamber containing a single *fmn* and of a chamber containing 13 *fmn* flies. **F.** With increased number of flies in a chamber (1, 2, 4, 8, versus 13), the *fmn* flies exhibited higher LIEs (ANOVA with *post hoc* Tukey; *F_4,55_* = 161.07, *p* < 0.0001; letters denote significant differences among groups; *n* = 12). **G.** In males, live same-sex but not opposite-sex flies substantially contribute to the loss of movement suppression. The single *fmn* male housed with 12 dummies, dead *CS* males or *CS* females showed no changes in inhibitory control; however, the single *fmn* male housed with 12 decapitated or 12 intact *CS* males displayed significantly greater LIEs (ANOVA with *post hoc* Tukey; *F_6,77_* = 189.96, *p* < 0.0001; *n* = 12). **H.** In females, either live same-sex or opposite-sex flies triggered impulsive flying in *fmn*. Like males, the single *fmn* female housed with 12 dummies or dead *CS* females showed robust inhibitory control; however, the single *fmn* female housed with either 12 decapitated, 12 intact *CS* females or 12 intact *CS* males displayed high LIE levels (ANOVA with *post hoc* Tukey; *F_6,77_* = 122.23, *p* < 0.0001; *n* = 12). **I.** Visual input is necessary for LIEs. The *fmn*’s LIEs was suppressed under infrared light (Student’s *t*-test; ***, *p* < 0.0001; *n* = 12). **J.** Olfactory input is important for LIEs. The *fmn* flies expressing tetanus toxin (TNT) in olfactory neurons (*Orco-GAL4>UAS-TNT*) showed normal movement suppression (ANOVA with *post hoc* Tukey; *F_2,33_* = 47.17, *p* < 0.0001; ***, *p* < 0.0001; *n* = 12). **K-L.** Gustatory input is crucial for LIEs. The *fmn* flies with TNT expressed in the major gustatory receptor-containing neurons (either *Gr32a-GAL4>UAS-TNT* or *Gr63a-GAL4>UAS-TNT*) displayed normal inhibitory control (For Figure 1K: ANOVA with *post hoc* Tukey; *F_2,33_* = 132.8, *p* < 0.0001; ***, *p* < 0.0001; *n* = 12; For Figure 1L: ANOVA with *post hoc* Tukey; *F_2,33_* = 122.24, *p* < 0.0001; ***, *p* < 0.0001; *n* = 12).

Dopamine is known to modulate inhibitory control and psychostimulant-induced hyper-dopaminergic states cause motor impulsivity ^51,52^. We examined the dopamine transporter (DAT) mutant *fumin* (*fmn*), which cannot reuptake released dopamine and thus has increased extracellular dopamine ^22,53^. When tested individually, *fmn* showed strong inhibitory control similar to wild-type *CS* (Figure 1D). However, when tested in a group, *fmn* showed significantly increased flying events (Figure 1E), with the frequency of these events rising as the number of flies in the chamber increased (Figure 1F). To further investigate, we exposed flies to wasp sound instead of strong airflow. Similar to the strong airflow condition, flies stopped moving upon exposure to wasp sound and maintained movement suppression when tested alone (Supplementary figures 1A and 1D) or in a group for *CS* (Supplementary figures 1B and 1C). However, when *fmn* flies were tested in a group, they displayed a high frequency of impulsive flying events, which we quantified as Loss of Inhibition Event (LIE; movements >60 mm/sec, Supplementary figures 1E and 1F). These findings collectively demonstrate that hyper-dopamine activity induces motor impulsivity only in a group setting.

### Social context influences impulsivity in *fmn* mutants

To identify the nature of a group setting that triggers impulsive flights, we tested a single *fmn* male or female with various objects or flies. Neither dummies nor dead flies had any effect. However, decapitated or intact live same-sex *CS* significantly provoked LIEs in both sexes (Figures 1G and 1H). In this mixed genotype population, *CS* flies were unaffected by the *fmn*’s LIEs and maintained movement suppression; however, the mere presence of live *CS* flies was sufficient to elicit LIEs in a *fmn* male or female. To assess the impact of opposite-sex context, we tested a *fmn* male in the presence of *CS* females and a *fmn* female in the presence of *CS* males. *CS* females did not induce LIEs in a *fmn* male (Figure 1G) however *CS* males triggered significant LIEs in a *fmn* female (Figure 1H). These findings validate social context as a critical factor interacting with hyper-dopamine to drive impulsivity and also reveal a sexual dimorphism in how *fmn* flies process opposite-sex social contexts.

Next we investigated which sensory modalities of social context are critical for LIEs. Groups of *fmn* flies were subjected to GNG under various sensory deprivation conditions: infrared light to impair vision ^54^ or silenced olfactory neurons for volatile pheromones (*Orco-GAL4>UAS-TNT*) ^55^ or silenced gustatory neurons detecting taste, pheromones and CO_2_ (*Gr32a-GAL4>UAS-TNT* and *Gr32a-GAL4>UAS-TNT*) ^34–36,56^. Deprivation of any single sensory modality completely suppressed LIEs in *fmn* (Figures 1I-L). This demonstrates that the entire sensory representation of other flies is essential to provoke impulsivity. Finally, we assessed whether mating status influenced LIEs. Both unmated and mated *fmn* flies exhibited similar LIEs in all tested conditions (Supplementary figures 2A and 2B), indicating that mating status is not a significant factor for impulsivity.

### Dopamine neurons projecting to the MB γ lobes drive impulsivity

The fly brain contains approximately 280 dopamine neurons with widespread projections ^57^. To identify the dopamine neurons involved in motor impulsivity, we performed rescue experiments. Restoring DAT expression in all dopamine neurons (*TH,DDC-GAL4>UAS-DAT*) or all neurons (*elav-GAL4>UAS-DAT*) fully suppressed the anomalous LIEs in *fmn* (Figure 2A). We hypothesized that ectopic DAT expression in postsynaptic sites proximal, but not distal, to the dopamine neurons driving motor impulsivity would rescue the LIE phenotype. Ectopic DAT expression in the MB (*OK107-GAL4>UAS-DAT*), a key brain structure for high-order functions ^58–60^, fully suppressed the LIE phenotype. DAT expression in other brain regions such as ellipsoid body (*c232-GAL4>UAS-DAT*), fan-shaped body (*c205-GAL4>UAS-DAT*), or neuropeptide F (*NPF-GAL4>UAS-DAT*) neurons had no significant effect (Figure 2A). These findings highlight the importance of the dopaminergic circuit involving the MB in regulating impulsivity.

**Figure 2.**
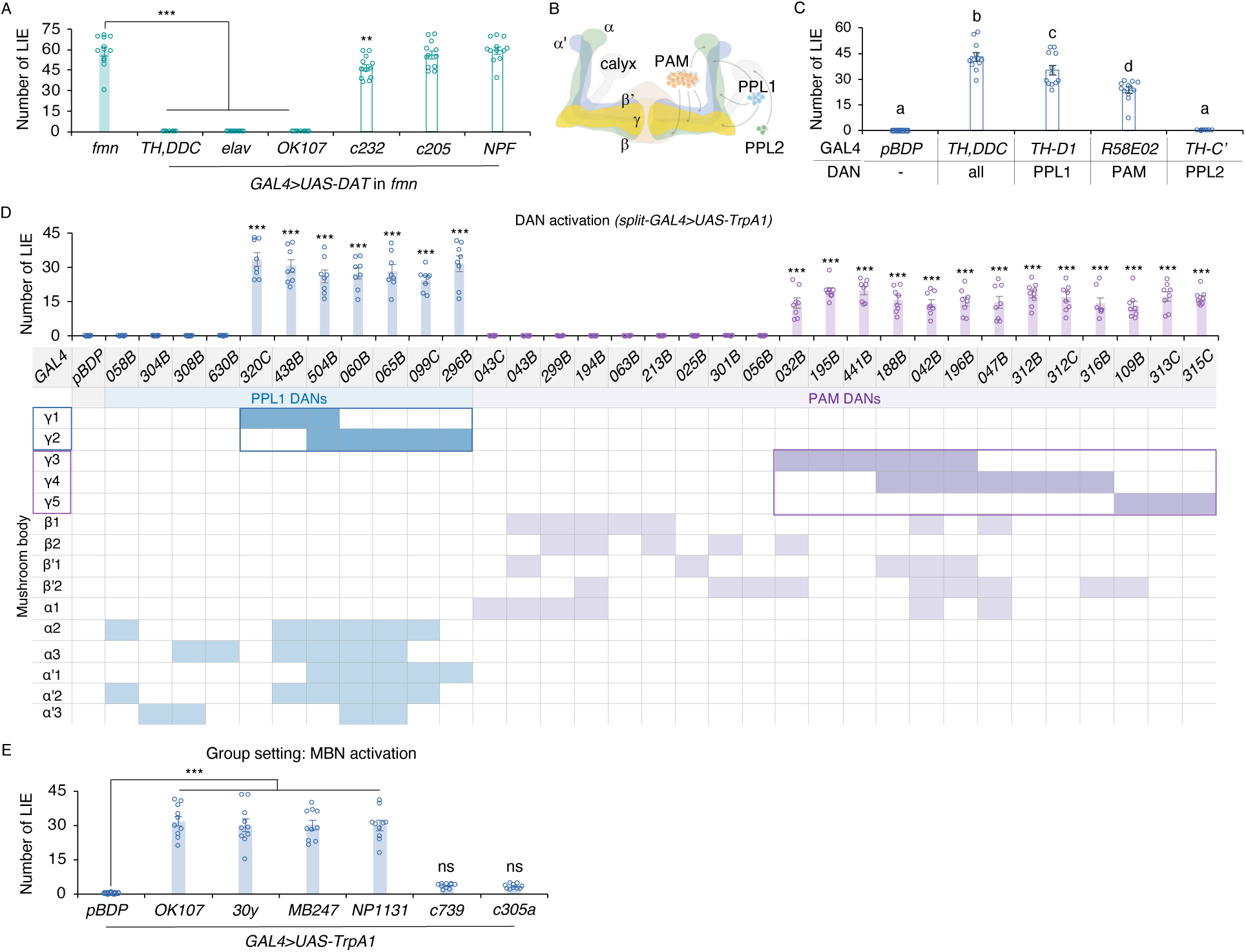
The dopamine neural circuit involving the MB lobes is important for impulsivity. **A.** DAT mutation rather than other factors is responsible for LIEs in *fmn*. The *fmn* flies carrying *UAS-DAT* and the GAL4 driver for dopamine neurons (*TH,DDC*), all neurons (*elav*) or all MB neurons (*OK107*), but not the GAL4 driver for ellipsoid body (*c232*), fan-shaped body (*c205*) or neuropeptide F (*NPF*), showed normal movement suppression (ANOVA with *post hoc* Dunnett’s; *F_6,77_* = 198.66, *p* < 0.0001; **, *p* < 0.001;***, *p* < 0.0001; *n* = 12), indicating that restored DAT expression in either dopamine or postsynaptic MB neurons rescues the *fmn*’s phenotype. Reinstated DAT expression in the ellipsoid body located close to the MB yielded marginal rescue. **B.** Scheme of three dopamine neuronal clusters (PAM, PPL1 and PPL2) projecting to the MB lobes (axons) and calyces (dendrites). The arrows demarcate the MB lobes (γ, αβ, α’β’). **C.** The dopamine neurons (DANs) important for inhibitory control. The flies with enhanced activity of all dopamine (*TH,DDC*), PPL1 (*TH-D1*) and PAM (*R58E02*) but not PPL2 (*TH-C’*) neurons showed significant LIEs (ANOVA with *post hoc* Tukey; *F_4,55_* = 121.15, *p* < 0.0001; *n* = 12). The *pBDP* containing promotor-less GAL4 (thus no GAL4 expression) was used as a control. **D.** Screening of the PPL1 and PAM DANs relevant for inhibitory control via ectopic activation in a group setting. To identify the specific PPL1 and/or PAM DANs important for inhibitory control, 33 split-GAL4 lines were used to drive *UAS*-*TrpA1* for ectopic activation of DANs. The *pDBP*-GAL4 driver served as control. Twenty out of the 33 split-GAL4 lines exhibited higher LIEs under the Go/No-Go test. Activation of PPL1 DANs (blue bars) having axonal projections to either the MB γ1 (*MB320C*, *MB438B*, or *MB504B*,) or γ2 (*MB504B*, *MB060B*, *MB065B*, *MB099C*, or *MB296B*) while activation of PAM DANs (green bars) having axonal projections to either the MB γ3 (*MB032B*, *MB195B*, *MB441B*, *MB188B*, *MB042B*, or *MB196B*), γ4 (*MB188B*, *MB042B*, *MB196B*, *MB047B*, *MB312B*, *MB312C*, or *MB316B*), or γ5 (*MB109B*, *MB313C*, or *MB315C*) resulted in augmented LIEs. The table specifies the PPL1 or PAM DAN projections to specific MB sub-compartments (ANOVA with *post hoc* Dunnett’s; *F_3,238_* = 43.91, *p* < 0.0001; ***, *p* < 0.0001; *n* = 8). **D.** In a group setting, activation of all MB neurons (*OK107* or *30Y*) or γ and αβ neurons (*MB247*) or γ (*NP1131*) led to high levels of LIE while activation of αβ (*c379*) or α’β’ (*c305a*) neurons caused minimal LIE increase (ANOVA with *post hoc* Dunnett’s; *F_6,63_* = 73.04, *p* < 0.0001; ns, *p* > 0.05; ***, *p* < 0.0001; *n* = 10).

Three dopamine neuronal clusters (PAM, PPL1 and PPL2) innervate the MB (Figure 2B) ^57^. To identify the specific dopamine neurons critical for motor impulsivity, we used GAL4 lines ^42,61^ to target each cluster: PPL1 (*TH-D1-GAL4*), PAM (*R58E02-GAL4*) and PPL2 (*TH-C’-GAL4*) cluster. By activating these clusters individually in a wild-type background, we assessed their contributions to LIEs. Activation of PPL1 (*TH-D1-GAL4>UAS-TrpA1*) or PAM (*R58E02-GAL4>UAS-TrpA1*) which innervate the MB lobes, induced significant LIEs. In contrast, activation of PPL2 (*TH-C’-GAL4>UAS-TrpA1*) innervating the MB calyces did not produce LIEs (Figure 2C). Notably, LIEs resulting from activation of individual clusters were less pronounced than those induced by simultaneous activation of all dopamine neurons. These results suggest that the combined PPL1 and PAM inputs to the MB lobes are critical for escalating dysfunctional inhibitory control and driving motor impulsivity.

The MB has three lobes (α/β, α’/β’, γ), each with well-defined subdomains such as the five subdomains in the γ lobe, which have distinct neural functions ^28,38,58–60^. The PPL1 and PAM DANs project axons to specific MB sub-compartments ^38^. To dissect the PPL1- and PAM-MB microcircuits involved in motor impulsivity, we used split-GAL4 lines targeting subsets of these DANs ^38^. Activation of PPL1 DANs innervating the MB γ1 (*MB320C-GAL4>UAS-TrpA1*, *MB438B-GAL4>UAS-TrpA1*, and *MB504B-GAL4>UAS-TrpA1*) or γ2 (*MB504B-GAL4>UAS-TrpA1*, *MB060B-GAL4>UAS-TrpA1*, *MB065B-GAL4>UAS-TrpA1*, *MB099C-GAL4>UAS-TrpA1*, and *MB296B-GAL4>UAS-TrpA1*) subdomains, as well as PAM DANs innervating γ3 (*MB032B-GAL4>UAS-TrpA1*, *MB195B-GAL4>UAS-TrpA1*, *MB441B-GAL4>UAS-TrpA1*, *MB188B-GAL4>UAS-TrpA1*, *MB042B-GAL4>UAS-TrpA1*, and *MB196B-GAL4>UAS-TrpA1*), γ4 (*MB188B-GAL4>UAS-TrpA1*, *MB042B-GAL4>UAS-TrpA1*, *MB196B-GAL4>UAS-TrpA1*, *MB047B-GAL4>UAS-TrpA1*, *MB312B-GAL4>UAS-TrpA1*, *MB312C-GAL4>UAS-TrpA1*, and *MB316B-GAL4>UAS-TrpA1*), or γ5 (*MB109B-GAL4>UAS-TrpA1*, *MB313C-GAL4>UAS-TrpA1*, and *MB315C-GAL4>UAS-TrpA1*) subdomains induced LIEs. In contrast, activation of PPL1 or PAM DANs innervating other MB regions did not trigger LIEs (Figure 2D). These findings indicate that PPL1 and PAM DANs mediate impulsivity via signaling to the MB γ lobes. Consistent with this, activation of MB γ neurons alone (*NP1131-GAL4>UAS-TrpA1*) was sufficient to induce LIEs in a group setting, even in the absence of hyper-dopamine. Activation of αβ (*c379-GAL4>UAS-TrpA1*) or α’β’ (*c305a-GAL4>UAS-TrpA1*) neurons did not produce LIEs. Moreover, activation of broader MB subsets (*MB247-GAL4>UAS-TrpA1*) or all MB neurons (*OK107-GAL4 or 30y-GAL4>UAS-TrpA1*) did not further increase LIEs (Figure 2E).

### Social Context and Hyper-Dopamine Converge in MB γ Lobes

As shown above, a group setting, but not a single-fly setting, triggered LIEs in *fmn* but not in *CS*, indicating that impulsivity requires both hyper-dopamine and social context. To identify where social context information converge with hyper-dopamine to drive impulsivity, we focused on the candidate neurons implicated in the process: PPL1 DANs, PAM DANs and MB γ neurons. We manipulated the candidate neurons in the absence of social context (i.e., a single-fly setting) and assessed LIEs. Activation of PPL1 (*TH-D1-GAL4>UAS-TrpA1*) or PAM (*R58E02-GAL4>UAS-TrpA1*) DANs did not induce LIEs (Figure 3A; Supplementary figure 3). Similarly, activating all DANs (*TH,DDC-GAL4>UAS-TrpA1*) or individual DAN subsets also failed to trigger LIEs (Figure 3A PPL2 (*TH-C’-GAL4>UAS-TrpA1*); Supplementary figure 3). These results suggest that DANs are not the site where dopamine and social context information converge to mediate impulsivity.

**Figure 3.**
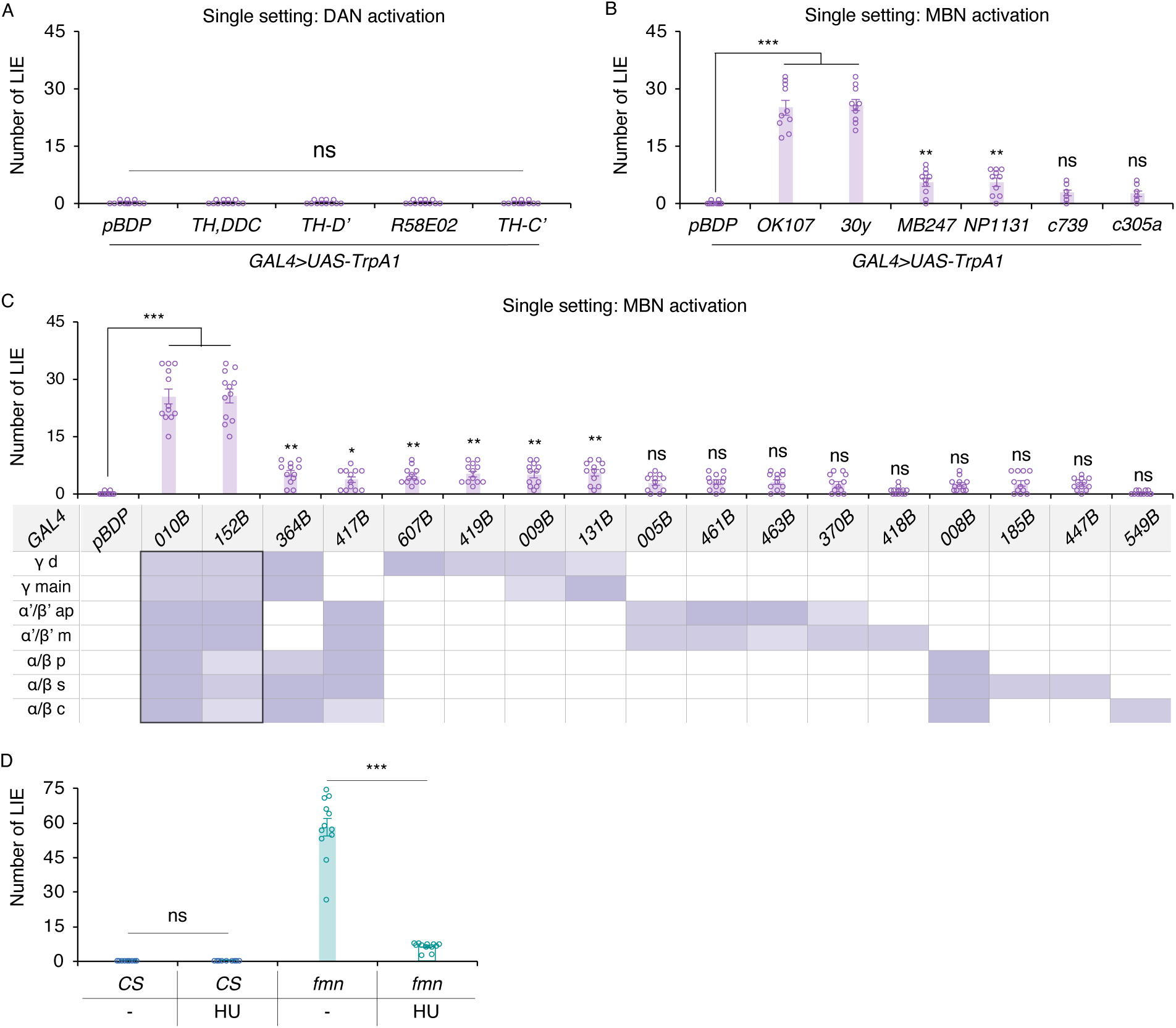
Hyper-dopamine signaling and social context information converge at the MB neurons for impulsivity. **A.** The single fly with enhanced activity in either all dopamine (*TH,DDC*), PPL1 (*TH-D1*), PAM (*R58E02*), or PPL2 (*TH-C’*) neurons did not show LIEs when tested alone similar to the promotor-less *pBDP*-GAL4 control (Kruskal-Wallis test; *p* = 0.990; *n* = 12). **B.** The single fly with all MB neuron activation (*OK107* or *30Y*) showed the substantial increase in LIEs while the fly with individual MB subset activation (*MB247* for γ and αβ neurons; *NP1131* for γ; *c739* for αβ; *c305a* for α’β’) displayed marginal increases. (ANOVA with *post hoc* Dunnett’s; *F_6,63_* = 92.90, *p* < 0.0001; ns, *p* > 0.05; **, *p* < 0.001; ***, *p* < 0.0001; *n* = 10). **C.** Screening of the MBNs relevant for inhibitory control via ectopic activation in a single setting. To identify the specific MBNs important for inhibitory control, 17 split-GAL4 lines were used to drive *UAS*-*TrpA1* for ectopic activation of either all MBNs or specific sub-compartments. The *pDBP*-GAL4 driver served as control. Activation of all MBNs (*MB010B* or *MB152B*; 2 out of the 17 split-GAL4 lines) exhibited markedly higher LIEs under the Go/No-Go test while activation of specific MB sub-compartments (15 out of 17 split-GAL4 lines) showed either marginal or no changes in LIE. The table specifies the expression patterns for the MBN sub-regions per each split-GAL4 line. (ANOVA with *post hoc* Dunnett’s; *F_17,198_* = 74.37, *p* < 0.0001; ns, *p* > 0.05; *, *p* < 0.05; **, *p* < 0.001; ***, *p* < 0.0001; *n* = 12). **D.** The *fmn*’s loss of inhibitory control requires functional MB. The MB-ablated *fmn* flies showed normal inhibition similar to *CS* flies (For *CS*: Student’s *t*-test; ns, *p* > 0.05; *n* = 12; For *fmn*: Student’s *t*-test; ***, *p* < 0.0001; *n* = 12).

Activation of MB γ (*NP1131-GAL4>UAS-TrpA1* for γ; *MB247-GAL4>UAS-TrpA1* for αβγ) resulted in small but significant increases in LIEs while activation of other MB subsets (*c739-GAL4>UAS-TrpA1* for αβ and *c305a-GAL4>UAS-TrpA1* for α’β’) had negligible effects (Figure 3B). Notably, all MB activation (*OK107-GAL4>UAS-TrpA1* or *30Y-GAL4> UAS-TrpA1*) produced substantial increases in LIEs (Figure 3B). These results were further supported by experiments using split-GAL4 lines that label specific MB neurons. Activation of all MB (*MB010B-GAL4* or *MB152B-GAL4*) caused a significant increase in LIEs, while activation of subsets such as γd, γ, γαβ, or αβa’β’ neurons produced small but still significant effects. Activation of all α’β’ or αβ neurons or their subsets had no effect (Figure 3C). Consistently, *fmn* flies with only γd–due to larval MB neuroblast ablation ^49,62^–exhibited normal inhibition in a group setting, behaving as if they were tested alone (Figure 3D). Together, these findings demonstrate that all MB lobes contribute to processing social context information, with the γ lobe serving as the key site where hyper-dopamine signals converge with social context information to drive motor impulsivity.

### Dopamine-D1 Receptor-cAMP Pathway Mediates Impulsivity

The inability to reuptake released dopamine in *fmn* flies likely leads to enhanced dopamine receptor signaling. To identify the dopamine receptor mediating motor impulsivity, we tested genetic interactions between *fmn* and *dumb*, *damb*, *der*, or *dd2r* with mutation in dDA1 (D1 dopamine receptor), DAMB (D5 dopamine receptor), DopEcR (Dopamine/Ecdysone receptor) or dD2R (D2 dopamine receptor), respectively ^23–26,63^. *dumb, damb* or *der* fully suppressed the LIEs in *fmn* in a group setting whereas *dd2r* had no effect (Figure 4A). Additionally, *fmn* flies with double-heterozygous *dumb*/*damb*, *dumb*/*der*, or *damb*/*der* displayed behavior similar to wild-type flies (Figure 4A). Consistent with these genetic results, pharmacological treatment with D1 receptor antagonist, but not D2 receptor antagonist, significantly reduced the LIEs in a dose-dependent manner (Supplementary figure 4). These results demonstrate that elevated dopamine signaling via the D1-like dopamine receptors mediates motor impulsivity.

**Figure 4.**
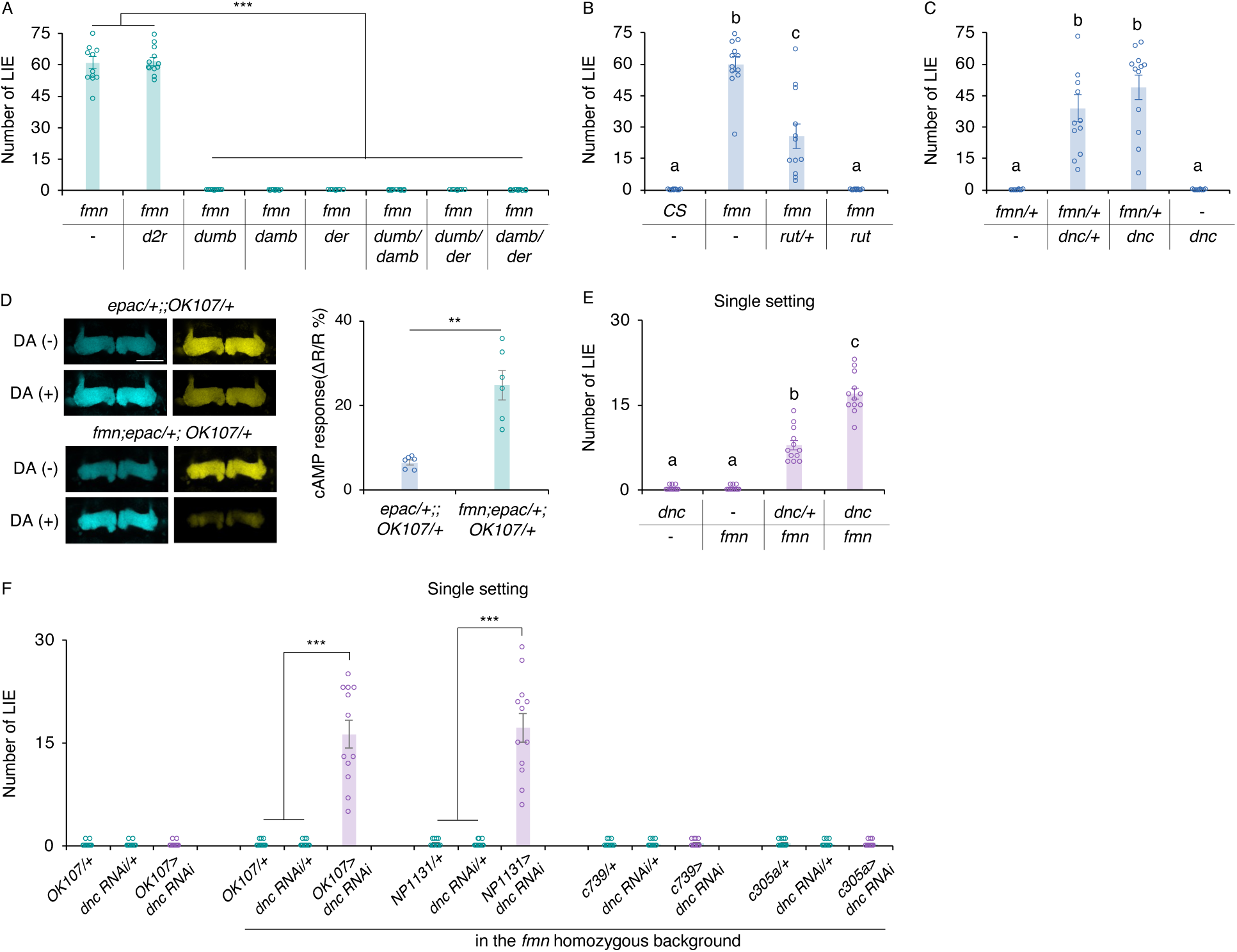
The D1 dopamine receptor dDA1–cAMP pathway in the MB is critical for the fly impulse control. **A**. The D1-like receptors are essential for inhibitory control. The *fmn* flies with mutation in either dDA1 (*dumb*), DAMB (*damb*) or DopEcR (*der*), but not with dD2R (*dd2r*), did not exhibit LIEs. Also, the *fmn* flies with the transheterozygous mutations in either dDA1 and DAMB (*dumb*/*damb*), dDA1 and DopEcR (*dumb*/*der*), or DAMB and DopEcR (*damb*/*der*) showed normal inhibitory control (ANOVA with *post hoc* Tukey; *F_7,88_* = 563.43, *p* < 0.0001; ***, *p* < 0.0001; *n* = 12). Genetic interaction between *fmn* and cAMP-associated genes *rutabaga* (*rut*; coding for adenylyl cyclase) and *dunce* (*dnc*; coding for phosphodiesterase). **B.** In a group setting, *fmn*’s LIEs was partially or fully suppressed by *rut/+* or *rut*, respectively (ANOVA with *post hoc* Tukey; *F_3,44_* = 68.50, *p* < 0.0001; *n* = 12). **C.** In a group setting, the *fmn/*+ and *dnc* flies exhibited robust inhibitory control while the *dnc/+;fmn/+* and *dnc;fmn/+* exhibited high LIEs (ANOVA with *post hoc* Tukey; *F_3,44_* = 34.2, *p* < 0.0001; *n* = 12). **D.** *fmn* displayed greater cAMP response. The control (*epac/+;;OK107 /+*) and *fmn* (*fmn;epac/+;OK107 /+*) brains expressing the cAMP sensor Epac1-camps (epac) in the MB were treated with 100 μM dopamine or HL3 (control). The dopamine-induced cAMP increase was higher in the γ lobe of *fmn* compared to the control (Student’s *t*-test; **, *p* < 0.001; *n* = 6). DA, dopamine; Scale bar, 50 μm. **E.** In a single setting, *dnc* interacts with *fmn* to cause dysfunctional inhibitory control. The *dnc/+;fmn*, and *dnc;fmn* exhibited higher LIEs compared to controls (ANOVA with *post hoc* Tukey; *F_3,44_* = 139.89, *p* < 0.0001; *n* = 12). **F.** In a single setting, *dnc* knockdown in either all MBNs or MB γ neurons in *fmn* led to the highly augmented LIEs compared to controls (Kruskal-Wallis and Mann-Whitney tests; ***, *p* < 0.0001; *n* = 12).

cAMP acts as a downstream signaling molecule for dDA1, DAMB and DopEcR *in vitro* ^25,64,65^. To investigate whether cAMP mediates motor impulsivity, we examined the genetic interaction of *fmn* with *rutabaga* (*rut*) defective in adenylyl cyclase that makes cAMP or *dunce* (*dnc*) defective in phosphodiesterase that degrades cAMP ^45,46^. As expected, *rut* suppressed *fmn*’s LIEs in a dose-dependent manner (*rut/+;fmn* and *rut;fmn*; Figure 4B). In contrast, negligible LIEs were observed in heterozygous *fmn* flies (*fmn/+*) while the addition of *dnc* to *fmn/+* significantly increased LIEs in a dose dependent manner (*dnc/+;fmn/+*, *dnc;fmn/+*; Figure 4C). However, *dnc* mutations without *fmn* did not induce LIEs in a group setting, indicating that cAMP elevation partly transmits *fmn*’s hyper-dopamine signal to drive impulsivity. To directly demonstrate that *fmn* elevates cAMP, we used the cAMP sensor Epac1-camps ^41^ and observed significantly larger cAMP increases in the MB γ lobe of *fmn* flies compared to controls (Figure 4D).

Next, we investigated whether cAMP is also involved in processing social context information. In a single-fly setting, *dnc* mutation alone had no effect on LIE like *fmn* (Figure 4E). However, when combined with *fmn,* the *dnc* mutation triggered LIEs in a dose-dependent manner (*dnc/+;fmn*, *dnc;fmn*; Figure 4E), indicating that cAMP is involved in social context processing. As shown previously, all three MB subsets are important for processing social context information. To determine whether cAMP is critical in these subsets, we knocked down *dnc* in all or individual MB subsets. In a wild-type background, *dnc* knockdown in all MB subsets did not affect behavior, similar to the genetic *dnc* mutant. However, *dnc* knockdown in all MB subsets in *fmn* led to a significant increase in LIEs (Figure 4F). Notably, *dnc* knockdown in the γ– but not in αβ or α’β’ –resulted in LIEs at levels comparable to those induced by *dnc* knockdown in all MB subsets. Together, these results demonstrate that hyper-dopamine and social context information converge onto cAMP signaling specifically in the MB γ lobe.

## Discussion

Organisms spend a significant portion of their lives in dynamic social settings. While prior works have examined how prior social experiences such as early life trauma, social isolation or social defeat shape inhibitory control ^66,67^, little attention has been given to how the immediate social context influences this critical cognitive function. This study provides new insights into the complex interplay between neurochemical signaling and social context in modulating impulsivity. Using the *Drosophila* model, we demonstrated that elevated dopamine levels alone are insufficient to cause impulsivity; instead, this trait emerges in the presence of social stimuli. Our findings identify the MB γ as a critical neural substrate where dopamine and social-context information converge, with the dopamine-D1 receptor-cAMP pathway playing a central role in this process.

Flies, though solitary, exhibit diverse social behaviors like courtship, aggression, collective action, and the formation of interaction networks ^68–71^. Our study identifies a previously unexplored interaction: the mere presence of same-sex peers in a shared environment disrupts inhibitory control and importantly, this effect is context-dependent—social context alone does not provoke impulsivity without a neural predisposition for increased dopamine signaling. A fly perceives other flies via multiple sensory modalities including visual and chemosensory inputs. Our results highlight the MB as the critical neural structure for integrating sensory and dopamine signals ^38,59,72^, underscoring its role as a hub for processing multimodal information and influencing behavioral outputs. Notably, the concurrent activation of all three MB lobes (γ, α/β, and α’/β’) is key to impulsivity. This suggests that simultaneous activation of the dopamine signaling and social context-processing circuits overwhelms the MB’s capacity for action selection, leading to dysregulated inhibitory control.

Our findings have significant implications for impulsivity research across species. Human and rodent studies of impulsivity typically occur in isolated conditions, omitting the potential influence of social context. This methodological limitation may partially explain the inconsistent associations reported between genetic variations and inhibitory control. While studies by Cornish et al. ^12^ and Cummins et al. ^13^, for example, demonstrate significant links between DAT1 gene (SLC6A3) variations and inhibitory control, other studies such as those by Kasparbauer et al. ^73^ and Gurvich and Rossell ^14^ report no such associations. These discrepancies might stem from social variables during task performance. Intriguingly, both Cummins et al. ^13^ and Kasparbauer et al. ^73^ have found significant correlations between DAT1 variants and fronto-striatal activity during response inhibition tasks. Our research demonstrates that social context is not merely a backdrop to behavior but an active modulator of cognitive processes like inhibitory control, underscoring the need to consider social variables when evaluating impulsivity and related cognitive processes. Impulsivity is a hallmark of various disorders including ADHD, autism spectrum disorder and substance use disorder, many of which are exacerbated in social settings^74–76^. Understanding the neural mechanisms that mediate social-context effects on behavior could inform interventions targeting the dynamic interplay between social stimuli and genetic or neural predispositions.

## Acknowledgments

We are grateful to the Bloomington Stock Center and Drs. Dubnau, Waddell, Wu, Anderson, Tomchik and Davis for sharing fly lines, the Cytometry, Screening and Imaging Core at Border Biomedical Research Center on confocal microscopy. We are also thankful to the past and current lab members for their help, discussion and support. This work was supported by the National Institutes of Health grant R21MH109953 (KAH), Brain & Behavior Research Foundation grant (KAH) and National Institutes of Health grant R16GM145548(KAH). National Institutes of Health grant R21MH109953 (KAH) Brain & Behavior Research Foundation grant (KAH) National Institutes of Health grant R16GM145548(KAH)

## Author contributions

Conceptualization: PRS, YCK, KAH

Methodology: PRS, YCK

Investigation: PRS, YCK, JMS

Visualization: PRS

Supervision: KAH

Writing—original draft: KAH

Writing—review & editing: PRS, YCK, JMS, KAH

## Data and materials availability

All raw data reported, and materials used in the study are available upon request from the corresponding authors.

## Supplemental information

**Supplementary figure 1.**
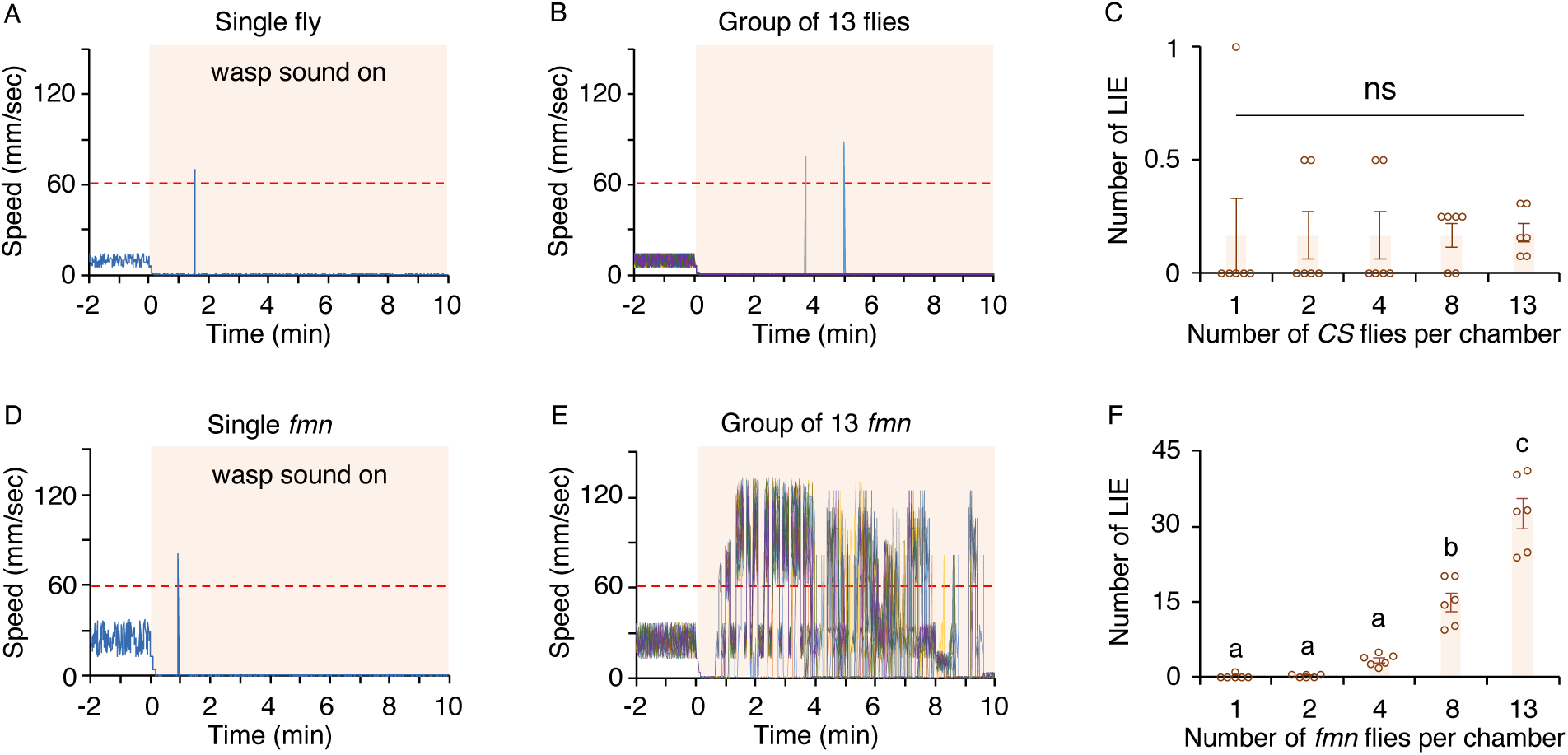
Wasp sound provokes movement suppression in *CS* and *fmn,* but the *fmn* flies are unable to maintain it. **A-B.** Representative movement traces of a chamber containing a single *CS* (**A**) and of a chamber containing 13 *CS* flies (**B**) under the GNG. The white area indicates Go phase where the wasp sound is off and the orange shaded area denotes No-Go phase where the wasp sound is on, and the traces exceeding the dotted red line on the 60 mm/sec mark represent a LIE. *CS* flies were moving in the chamber during the Go phase and upon exposure to wasp sound exhibited movement suppression (**A-B**). **C.** The number of *CS* flies in chamber does not impact impulsivity trigerred by wasp sound. (1, 2, 4, 8, versus 13; Kruskal-Wallis test, *p* = 0.571, *n* = 6). *fmn* show augmented impulsivity under the GNG when tested in a group. **D-E.** Representative movement traces of a chamber containing a single *fmn* and of a chamber containing 13 *fmn* flies. **F.** With increased number of flies in a chamber (1, 2, 4, 8, versus 13), the *fmn* flies exhibited elevated LIEs (ANOVA with *post hoc* Tukey; *F_4,25_* = 74.55, *p* < 0.0001; *n* = 6).

**Supplementary figure 2.**
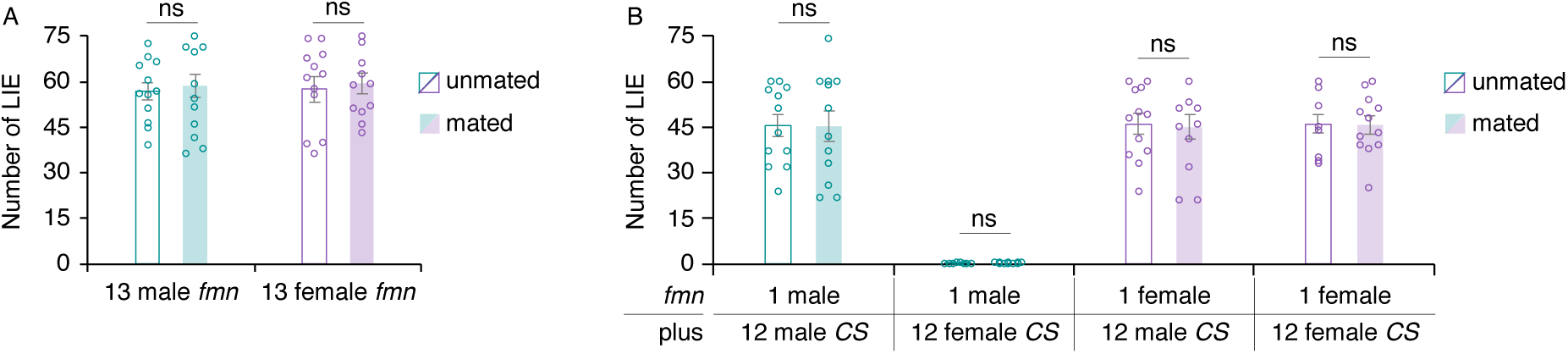
The social context-dependent impulsivity is independent mating status. **A.** The mating status of the group-tested *fmn* flies does not influence impulsivity. In both sexes, the unmated versus the mated *fmn* do not differ in LIEs (For male: Student’s *t*-test; ns, *p* > 0.05; *n* = 12; For female: Student’s *t*-test; ns, *p* > 0.05; *n* = 12). **B.** In either sex, the mated and unmated *fmn* displayed comparable LIEs across all social contexts tested. ( For 1 *fmn* male with 12 *CS* male: Student’s *t*-test; ns, *p* > 0.05; *n* = 12; For 1 *fmn* male with 12 *CS* female: Student’s *t*-test; ns, *p* > 0.05; *n* = 12; For 1 *fmn* female with 12 *CS* male: Student’s *t*-test; ns, *p* > 0.05; *n* = 12; For 1 *fmn* female with 12 *CS* female: Student’s *t*-test; ns, *p* > 0.05; *n* = 12.

**Supplementary figure 3.**
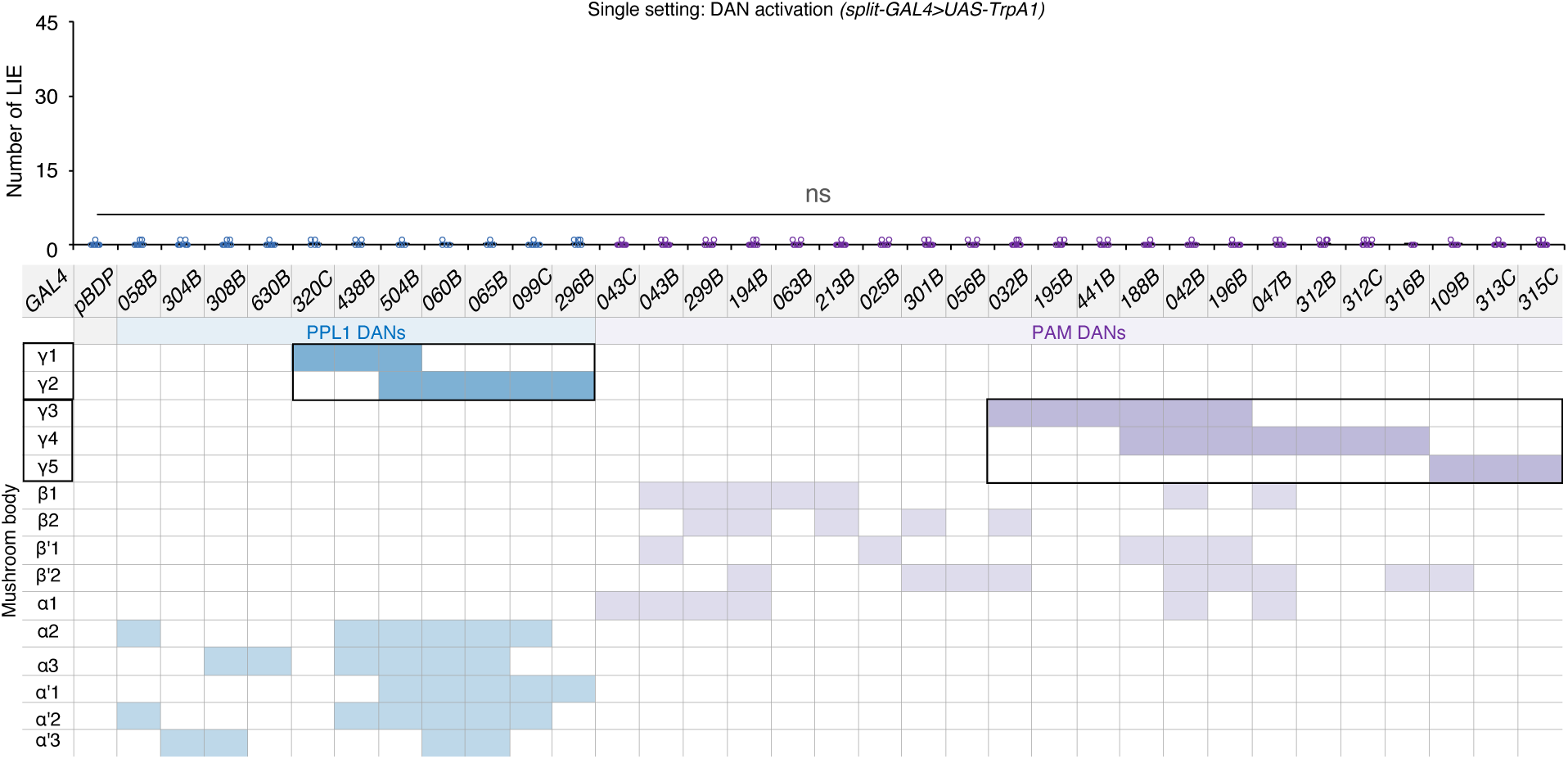
Screening of the PPL1 and PAM DANs relevant for inhibitory control via ectopic activation in a single setting. When ectopically activated, none of the 33 split-GAL4 lines expressing *UAS*-*TrpA1* displayed LIEs different from control. (Kruskal-Wallis test; *p* = 1.000; *n* = 8)

**Supplementary figure 4.**
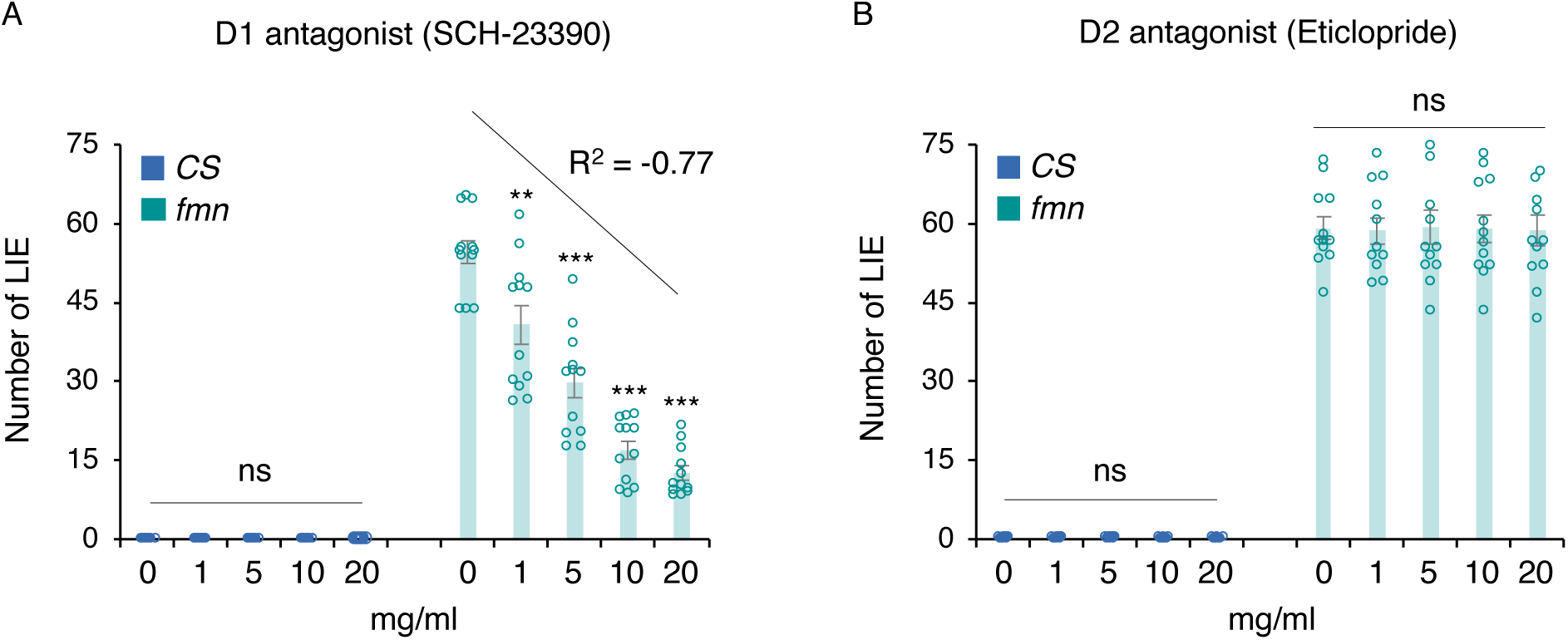
D1 but not D2 antagonist treatments inhibit the *fmn*’s LIEs. *CS* and *fmn* flies were treated with various concentrations of either D1 receptor antagonist (SCH-23390; **A**) or D2 receptor antagonist (Eticlopride; **B**). D1 but not D2 antagonist-fed *fmn* flies displayed dose-dependent decreases in LIEs. (For supplementary figure 4A, *CS*: ANOVA; *F_4,55_* = 1.05, *p* = 0.392; For supplementary figure 4A, *fmn*: ANOVA with *post hoc* Dunnett’s; *F_4,55_* = 47.77, *p* < 0.0001; **, *p* < 0.01; ***, *p* < 0.001; n = 12; For supplementary figure 4B, *CS*: ANOVA; *F_4,55_* = 1.82, *p* = 0.139; For supplementary figure 4B, *fmn*: ANOVA; *F_4,55_* = 0.01, *p* = 1.000; *n* = 12).

